# Prediction of peptide hormones using an ensemble of machine learning and similarity-based methods

**DOI:** 10.1101/2023.05.15.540764

**Authors:** Dashleen Kaur, Akanksha Arora, Palani Vigneshwar, Gajendra P.S. Raghava

**Author notes:** **Corresponding author** Prof. Gajendra P. S. Raghava, Head and Professor, Department of Computational Biology, Indraprastha Institute of Information Technology, Delhi, Okhla Industrial Estate, Phase III, (Near Govind Puri Metro Station), New Delhi, India – 110020 Office: A-302 (R&D Block), Website: http://webs.iiitd.edu.in/raghava/. Equal Contribution (Dashleen Kaur, Akanksha Arora). **Authors’ Information** Dashleen Kaur Akanksha Arora Palani Vigneshwar Prof. Gajendra P. S. Raghava. **Author’s Biography**. 1. Dashleen Kaur is currently pursuing an M. Tech. in Computational Biology at Department of Computational Biology, Indraprastha Institute of Information Technology, New Delhi, India. 2. Akanksha Arora is currently pursuing a Ph.D. in Computational Biology at Department of Computational Biology, Indraprastha Institute of Information Technology, New Delhi, India. 3. Palani Vigneshwar is currently pursuing an M. Tech. in Computer Science with specialization in Artificial Intelligence (CSAI) at Department of Computer Science and Engineering, Indraprastha Institute of Information Technology, New Delhi, India. 4. Gajendra P. S. Raghava is currently working as a Professor and Head of Department of Computational Biology, Indraprastha Institute of Information Technology, New Delhi, India.

## Abstract

Peptide hormones are genome-encoded signal transduction molecules released in multicellular organisms. The dysregulation of hormone release can cause multiple health problems and it is crucial to study these hormones for therapeutic purposes. To help the research community working in this field, we developed a prediction server that classifies hormonal peptides and non-hormonal peptides. The dataset used in this study was collected for both plants and animals from Hmrbase2 and PeptideAtlas databases. It comprises non-redundant 1174 hormonal and 1174 non-hormonal peptide sequences which were combined and divided into 80% training and 20% validation sets. We extracted a wide variety of compositional features from these sequences to develop various Machine Learning (ML) and Deep Learning (DL) models. The best performing model was logistic regression model trained on top 50 features which achieved an AUROC of 0.93. To enhance the performance of ML model, we applied Basic Local Alignment Search Tool (BLAST) to identify hormonal sequences using similarity among them, and motif search using Motif-Emerging and Classes-Identification (MERCI) to detect motifs present in hormonal and non-hormonal sequences. We combined our best performing classification model, i.e., logistic regression model with BLAST and MERCI to form a hybrid model that can predict hormonal peptide sequences accurately. The hybrid model is able to achieve an AUROC of 0.96, an accuracy of 89.79%, and an MCC of 0.8 on the validation set. This hybrid model has been incorporated on the publicly available website of HOPPred at https://webs.iiitd.edu.in/raghava/hoppred/.

## Introduction

The peptide hormones are a varied group of genome-encoded regulatory molecules with a specialized and important purpose of transferring specific information between cells and organs. This sort of molecular communication emerged and developed into a sophisticated system for regulating growth, development, and homeostasis in the early stages of evolution [1]. A wide range of reasons, including metabolite deposition in cells, surgical removal of glands or destruction of glands, can cause inadequate production of hormones. A frequent cause of hormone reduction is autoimmunity, and less frequently, it is also caused by genetic anomalies. Altered levels of hormones can disrupt the delicate equilibrium of the body and lead to endocrine disorders like endocrine neoplasia and diabetes [2]. It is thus indispensable to focus on the therapeutics of endocrine disorders associated with hormonal imbalances.

Peptide hormones, like all other peptides, are effective targeted signaling molecules that attach to specific cell surface receptors and cause intracellular effects. They are a great place to start while developing novel treatments because of their appealing pharmacological profile, target specificity and inherent qualities. It was, in fact, the foundational studies into natural hormones like insulin and gonadotropin-releasing hormone (GnRH), which marked the beginning of research into therapeutic peptides [3]. Their exceptional tolerability and efficacy have been demonstrated to be a direct result of their specificity. This feature may also distinguish between peptides and conventional small molecules. The depiction of mechanism of peptide hormones has been shown in Figure 1.

**Figure 1:**
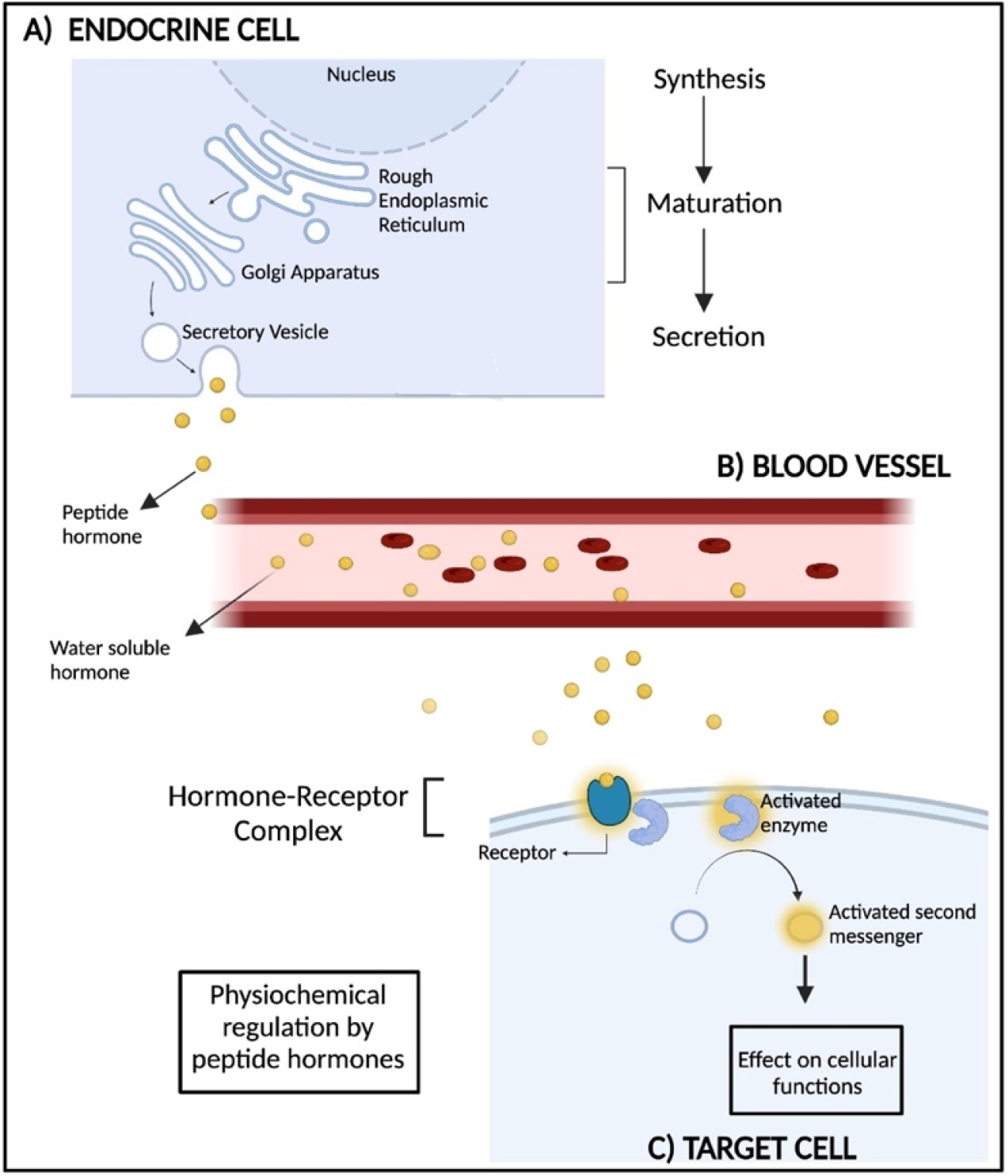
Mechanism of action of peptide hormones

Several peptide medications act as “replacement therapies,” restoring or supplanting peptide hormones when endogenous levels are insufficient or absent. They are intrinsic signalling molecules for a variety of physiological functions, which opens the possibility for therapeutic intervention emulating natural routes [4]. One such example is insulin which has helped thousands of people with diabetes since its introduction as the first peptide medication for commercial use in 1923 [3]. While replicating peptides generated from nature was formerly the discovery and development trend in the therapeutic peptide sector, it has recently changed to the rational design of peptides with desirable physiological and biochemical properties. Peptide-based natural hormone analogues have been developed as drug candidates. Given their enormous therapeutic potential, it can be anticipated that therapeutic hormone peptides will continue to draw research attention and see long-term success.

Several methods have been proposed in the past for predicting and designing therapeutic peptides. TPpred-ATMV improves the prediction of therapeutic peptides, THPep predicts the tumour-homing peptide (THP), AntiCP 2.0 for the prediction of anti-cancer peptides, and PrMFTP has been developed to predict multi-functional therapeutic peptides [5–8]. As far as we know, there is no method created particularly to predict peptide hormones. In this study, we have proposed an approach HOPPred to classify the peptide hormone and non-hormone peptide sequences. The complete architecture of HOPPred is given in Figure 2.

**Figure 2.**
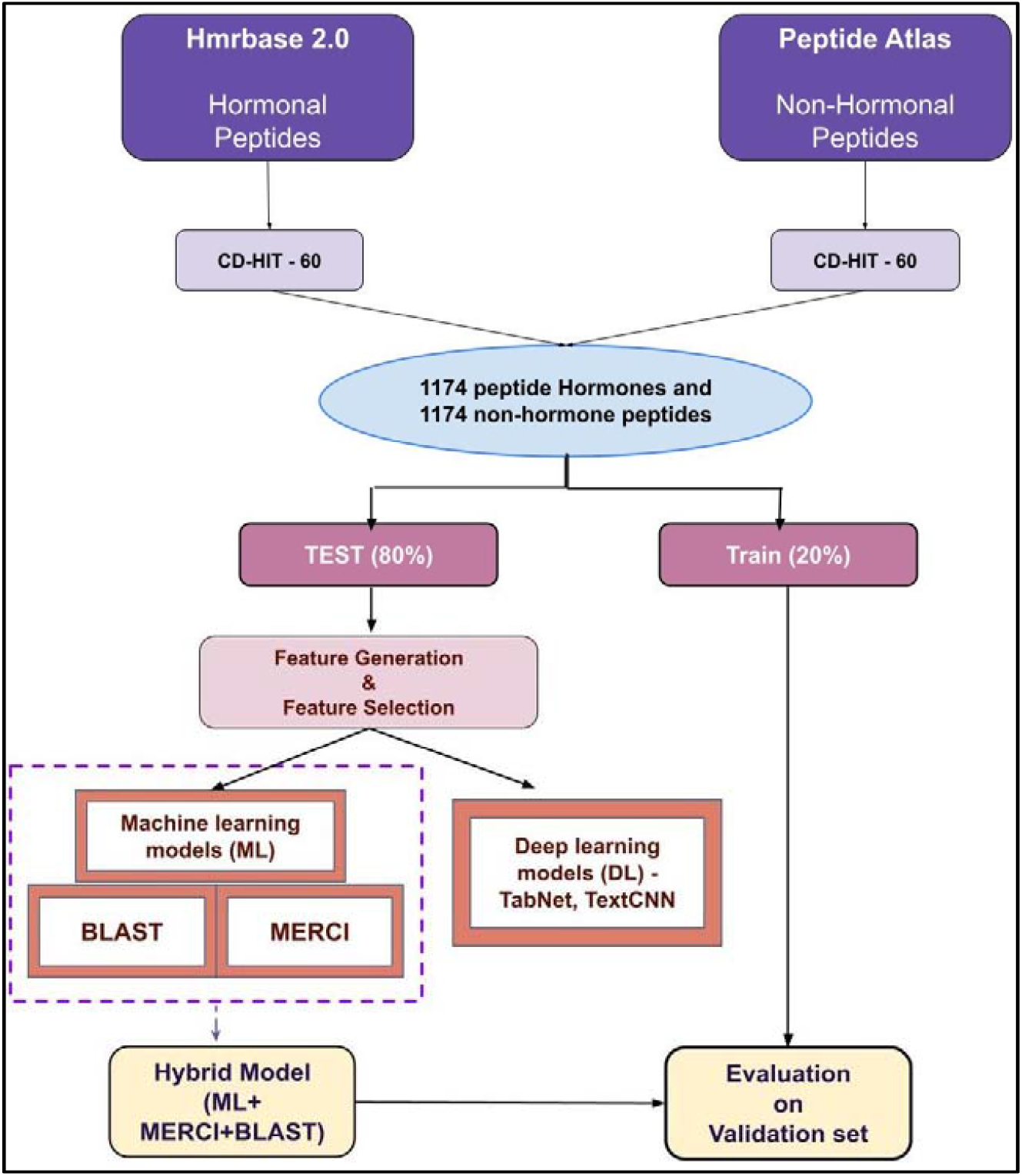
Architecture of HOPPred

## Methods

### Dataset compilation and preprocessing

We took 5729 peptide hormone sequences for both plants and animals from the Hmrbase2 database [9]. These are mature hormone sequences devoid of the signal and precursor areas. All duplicate peptides, ones that were longer than 41 and those that were shorter than 11 amino acids, were eliminated as were no peptides shorter than 11 amino acids in the data and a very few peptides with the length more than 41. Redundant peptides were removed and CD-HIT 60% was applied within the positive and negative datasets to make sure there are no peptides with more than 60% similarity [10]. Ultimately, we obtained 1174 peptide hormones, which we refer to in this work as the positive dataset. For the non-hormone peptide dataset, i.e., the negative dataset, we used PeptideAtlas database [11]. About 1174 non-hormone peptides were selected randomly from PeptideAtlas, and amino acid composition for these peptides were calculated to assess if the composition matches the standard amino acid composition of proteins/peptides present in nature.

### Feature generation and selection

Structural and functional annotations of peptide sequences were performed using Pfeature. It computes the descriptors of a sequence and gives a vector of 9149 features [12]. Using Pfeature we calculated all 9149 features, which included 15 categories of characteristics and descriptors, including amino acid compositions, tripeptide compositions, dipeptide compositions etc.

It is important to determine the few significant features out of the plethora of vast number of features [13]. To do that, we utilised the Python programming language’s Scikit-learn package and the Recursive Feature Elimination (RFE) technique, with Logistic Regression serving as the estimator [14]. The features were chosen from the standardized data obtained using the StandardScaler from Scikit-learn package [14]. Until a certain number of features have been attained, this feature selection approach recursively deletes the weakest features from the set. Unlike the other feature selection methods, which focus on individual properties of features, RFE focuses on features which affect the performance of the model [15]. We identified and used the top 50 most relevant features in the machine learning models which have been described in Supplementary Table S1.

### Machine Learning Techniques

We classified peptide hormones and non-hormonal peptides using a variety of machine-learning algorithms like Random Forest (RF), Decision Tree (DT), K-Nearest Neighbors (KNN), Gaussian Naïve Bayes (GNB), Logistic Regression (LR), and XGBoost (XGB) classifiers, using a python-based toolkit called Scikit-learn. DT classifier is a non-parametric supervised machine learning model which identifies the output by learning decision rules from input, ensemble-based RF classifier trains several decision trees to prodigy a single tree, LR classifier calculates the likelihood of an event using logistic function, KNN classifier bases its forecast on the highest number of votes cast in favour of the class that is closest to the nearest neighboring data point, GNB classifier is a probabilistic classifier based on Bayes’ theorem, and XGB classifier is a distributed gradient-boosted decision tree machine learning library that avails parallel tree boosting [16–21].

### Deep Learning Techniques

#### Tabnet

We applied TabNet on a set of 9149 biological features which were extracted using Pfeature. TabNet is a deep learning model applied on a tabular data. At each stage of the model, it chooses which features to use by giving sequential attention. This facilitates interpretability and improved learning as the most useful features are used. Tabnet utilizes a single deep learning architecture for both feature selection and reasoning known as soft feature selection [22].

#### TextCNN

We also used TextCNN method where we considered sequences as text and classified them using convolution neural networks. The padding of sequences was done to make the lengths same for all sequences. Then, we took the word length of 3 alphabets and divided the sequences into set of words. We then applied TextCNN to the matrix of 8000 features (20*20*20 = 8000), where, 20 is the possible amino acids that can be present at a single place [23].

### Five-fold cross-validation

To prevent bias and overfitting in the derived models, we used a 5-fold cross-validation procedure. The entire data was split into training data and validation data in the 80 to 20 ratio. The machine learning models were evaluated using the 5-fold cross-validation technique on 80% of training data. The training data is split into five folds, utilising four folds for training and the remaining fold as a test set for internal validation. Five repetitions are run through this process to give each fold a chance to be the test fold. The remaining 20% data is kept aside for external validation of the model. Many approaches have used this technique for the validation of data [24].

### Evaluation of parameters

The evaluation of the prediction model is one of the most important processes. Several parameters, which may be threshold-dependent or threshold-independent, can be used to evaluate the prediction models. Threshold-dependent parameters are sensitivity which measures how well the model can predict the hormones (Equation 1); specificity for the correct prediction of non-hormones (Equation 2), accuracy which shows the proportion of hormones and non-hormones that were successfully predicted (Equation 3); and Matthew’s correlation coefficient (MCC) shows the relationship between predicted and observed values (Equation 4). The threshold-independent parameter, the area under the receiver operating characteristic curve (AUC) between the sensitivity and 1-specificity, tells the capacity of the model to differentiate between the classes.

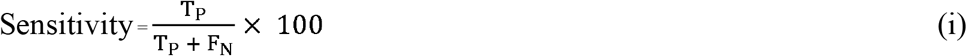

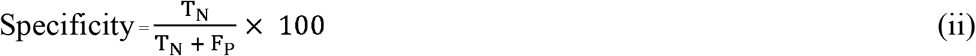

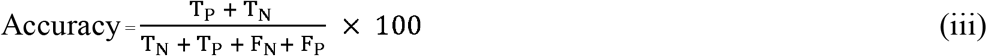

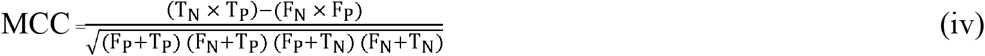

Where, T_P_, T_N_, F_P_ and F_N_ stand for true positive, true negative, false positive and false negative, respectively.

### BLAST for similarity search

It is common practice to annotate protein and nucleotide sequences using the Basic Local Alignment Search Tool (BLAST) [25]. We used BLAST shortp used for peptide sequences to identify hormonal sequences based on the similarity between sequences.

For this, we created a database of hormonal and non-hormonal sequences, and the query sequences were searched against this database. Peptide hormones were identified using the top hit at various E-value cutoffs. For eg., if a query sequence was hit against the hormonal peptide sequence, it was classified as hormone otherwise it was classified as non-hormone. Several researchers have used and thoroughly annotated this methodology [24,26]. The BLAST database was created using all sequences present in the training dataset. For evaluating the training set sequences, we hit all the sequences against the database and took the first hit after self-hit into account whereas for validation set sequences, we used the first hit to determine whether a query sequence is hormonal or non-hormonal.

### Motif analysis

Motif-EmeRging and Classes-Identification (MERCI) tool is used to find motifs in the set of peptide sequences [27]. The information provided by motif analysis relates to recurring patterns seen in peptide hormone sequences. The software utilises Perl script with the default settings to find motifs in the files. This software helps to discover novel motifs and carry out various motif-based analyses. It takes FASTA file of the sequences at input and gives out a set of repeating sequence patterns called as motifs and their occurrence positions in the sequences. In this study, we have found out the motifs that exclusively occur in hormonal and non-hormonal sequences.

### Hybrid approach

To improve the model’s prediction in this study, a hybrid strategy has also been used. The hybrid approach is a weighted scoring method that combines three different techniques to calculate the score: an approach based on similarity using BLAST, an approach based on motif using MERCI, and an approach using machine learning (ML). First, BLAST was used to categorise the provided protein sequence with an E-value of 10^−1^. We gave the positive predictions (peptide hormones) a weight of “+0.5,” the negative predictions (non-hormone peptides) a weight of “-0.5,” and no hits a weight of “0.” Second, MERCI was used to categorise the same peptide sequence. When the motifs were discovered, we gave them a score of “+0.5,” and when they were not, we gave them a score of “0.” Third, ML techniques were used to classify hormonal and non-hormonal peptides. If a peptide was classified as hormonal, we gave it a score of “+0.5” and “-0.5” if it was identified as non-hormonal. The scoring system is explained in detail in equations 1 and 2. In the hybrid technique, the total score was computed by combining the results of all three approaches. The peptide sequence is classified as hormonal or non-hormonal on the overall hybrid score (M’’) as defined in equation 2. The overall hybrid score varies from the range -1 to 2.

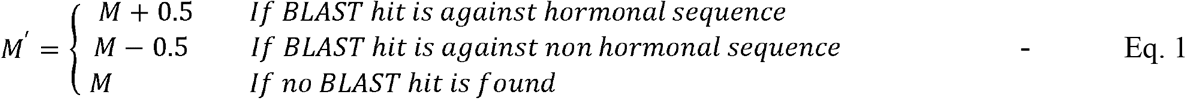

Here, *M* = Prediction score obtained from ML-based approach and M′ = Score obtained after adding scores from BLAST-based approach

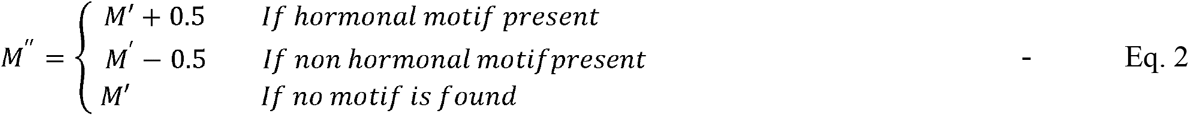

Here, M″= Hybrid score obtained from ML-based approach, BLAST-based approach, and motif-based approach ranging from -1 to 2

## Results

### Analysis of Amino Acid Composition

We performed the amino acid composition analysis for hormonal and non-hormonal peptides to find out the difference between two classes. Figure 3 depicts the amino acid composition of peptide hormones and non-hormonal peptides along with the p-values for each amino acid, making the compositional difference between them quite evident. We performed a two-sided Mann-Whitney U test to compare amino acid composition values of two groups, i.e., hormonal peptides and non-hormonal peptides. We found that the composition of cysteine, aspartic acid, phenylalanine, glycine, asparagine, proline, arginine, serine, and tyrosine were significantly increased in hormonal peptide sequences than non-hormonal peptide sequences with p-values<0.05. The composition of amino acids like glutamic acid, isoleucine, lysine, leucine, methionine, glutamine, threonine, and valine were significantly decreased in hormonal peptide sequences than non-hormonal peptide sequences with p-values<0.05.

**Figure 3:**
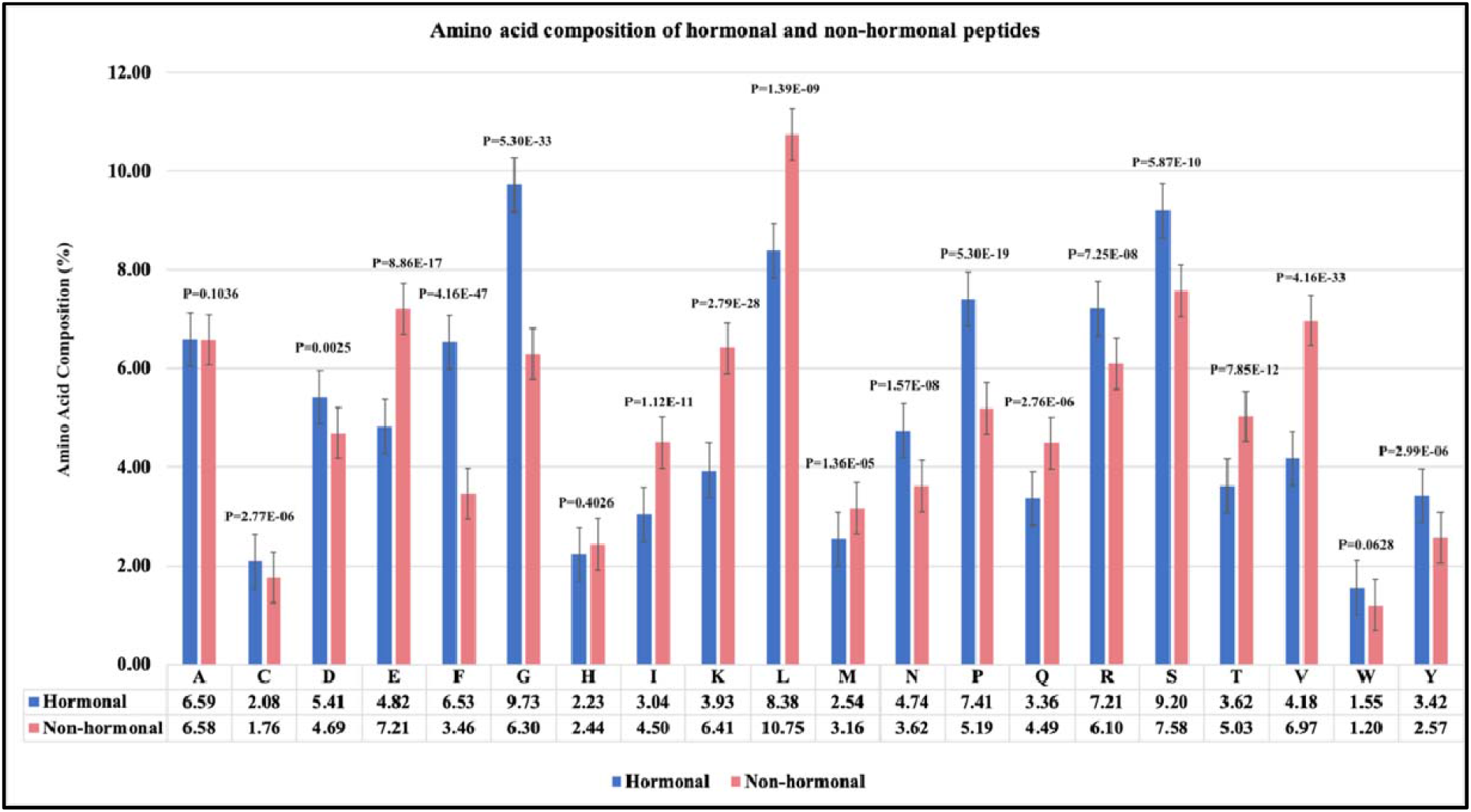
Amino acid composition analysis for peptide hormones and non-hormone peptides

### Performance of ML-Based Techniques on Selected Features

First, we computed a variety of features using the Pfeature software and removed the unnecessary features using the RFE feature selection methodology as mentioned in the Materials and Methods section. For comparison purposes, we developed multiple models based on the top 30 and top 50 features in the dataset and assessed them on the training and validation datasets. All the machine learning methods were used on the datasets with amino acid composition (AAC), top 30 and top 50 features to create peptide hormone prediction models. Hyperparameter tuning was used to optimize parameters, and the best model employed both the web server and standalone software.

The set of top 50 features performed best and were chosen as the final features in our prediction model, as evidenced by the results - LR (training: 0.94 AUROC and validation: 0.93 AUROC) is the best model, followed by RF (training: 0.91 AUROC and validation: 0.90 AUROC). The performance of various models employed on AAC, top 30 and top 50 features have been summarised in Table 1.

**Table 1:** Results for Machine Learning (ML) models developed for Amino Acid Composition (AAC), top 30 and top 50 features selected using RFE.

| Machine Learning Methods |  |  |  |  |  |  |  |  |  |  |
| --- | --- | --- | --- | --- | --- | --- | --- | --- | --- | --- |
| Amino Acid Composition (AAC) |  |  |  |  |  |  |  |  |  |  |
| Model | Training |  |  |  |  | Validation |  |  |  |  |
|  | Sens (%) | Spec (%) | Acc (%) | AUROC | MCC | Sens (%) | Spec (%) | Acc (%) | AUROC | MCC |
| **RF** | **80.79** | **80.34** | **80.56** | **0.89** | **0.61** | **79.31** | **84.03** | **81.70** | **0.89** | **0.63** |
| **LR** | 72.19 | 73.08 | 72.63 | 0.81 | 0.45 | 74.57 | 70.17 | 72.34 | 0.80 | 0.45 |
| **XGB** | 80.68 | 80.23 | 80.46 | 0.88 | 0.61 | 77.59 | 81.93 | 79.79 | 0.87 | 0.60 |
| **KNN** | 62.10 | 63.25 | 62.67 | 0.68 | 0.25 | 71.12 | 63.02 | 67.02 | 0.72 | 0.34 |
| **GNB** | 71.231 | 70.62 | 70.93 | 0.79 | 0.42 | 75.43 | 71.43 | 73.40 | 0.79 | 0.47 |
| **DT** | 66.67 | 67.20 | 66.93 | 0.72 | 0.34 | 60.78 | 66.39 | 63.62 | 0.70 | 0.27 |
| **SVC** | 74.10 | 72.33 | 73.22 | 0.81 | 0.46 | 76.72 | 68.49 | 72.55 | 0.79 | 0.45 |
| Top 30 features |  |  |  |  |  |  |  |  |  |  |
| Model | Training |  |  |  |  | Validation |  |  |  |  |
|  | Sens (%) | Spec (%) | Acc (%) | AUC | MCC | Sens (%) | Spec (%) | Acc (%) | AUC | MCC |
| **RF** | 80.68 | 80.77 | 80.72 | 0.90 | 0.61 | 77.59 | 83.61 | 80.64 | 0.88 | 0.61 |
| **LR** | **85.99** | **86.11** | **86.05** | **0.92** | **0.72** | **84.05** | **83.61** | **83.83** | **0.90** | **0.68** |
| **XGB** | 81.10 | 81.52 | 81.31 | 0.90 | 0.63 | 80.17 | 80.25 | 80.21 | 0.89 | 0.60 |
| **KNN** | 77.81 | 76.60 | 77.21 | 0.86 | 0.54 | 76.72 | 76.47 | 76.60 | 0.84 | 0.53 |
| **GNB** | 65.39 | 66.45 | 65.92 | 0.74 | 0.32 | 78.45 | 56.30 | 67.23 | 0.75 | 0.36 |
| **Top 50 features** |  |  |  |  |  |  |  |  |  |  |
| **Model** | **Training** |  |  |  |  | **Validation** |  |  |  |  |
|  | **Sens (%)** | **Spec (%)** | **Acc (%)** | **AUC** | **MCC** | **Sens (%)** | **Spec (%)** | **Acc (%)** | **AUC** | **MCC** |
| **RF** | 84.29 | 84.08 | 84.18 | 0.91 | 0.68 | 80.17 | 84.03 | 82.13 | 0.90 | 0.64 |
| **LR** | **87.05** | **87.40** | **87.22** | **0.94** | **0.74** | **85.34** | **86.55** | **85.96** | **0.93** | **0.72** |
| **XGB** | 84.18 | 84.29 | 84.24 | 0.92 | 0.68 | 77.59 | 84.45 | 81.06 | 0.90 | 0.62 |
| **KNN** | 56.37 | 46.79 | 51.60 | 0.52 | 0.03 | 57.33 | 47.90 | 52.55 | 0.56 | 0.05 |
| **GNB** | 24.52 | 95.94 | 60.12 | 0.76 | 0.29 | 22.84 | 97.06 | 60.43 | 0.83 | 0.30 |
| **DT** | 72.72 | 72.11 | 72.42 | 0.78 | 0.45 | 72.41 | 73.95 | 73.19 | 0.79 | 0.46 |

### Performance Deep Learning Techniques

#### TextCNN

TextCNN is a deep learning method to perform text classification tasks [23]. We divided the sequences into the words of length = 3 and used them to build our deep learning model. This model was able to achieve an AUROC of 0.98 on the training dataset, however, it was not able to perform well on the testing dataset giving an AUROC of 0.90. The MCC was also seen to decrease from 0.88 in training dataset to 0.67 in testing dataset.

#### TabNet

TabNet is a deep learning method that uses tabular data as input and is trained using gradient-descent based optimization [22]. The 9149 features generated from the Pfeature were passed as features through the TabNet model to get the train AUROC as 0.81 and the validation AUROC as 0.75 with an MCC of 0.61 for training dataset and 0.57 for validation dataset. The performances of the deep learning models – TextCNN and TabNet have been summarised in Table 2.

**Table 2:** The performance of deep learning (DL) models (TextCNN and TabNet)

| Deep Learning Methods |  |  |  |  |  |  |  |  |  |  |
| --- | --- | --- | --- | --- | --- | --- | --- | --- | --- | --- |
| Model | Training |  |  |  |  | Validation |  |  |  |  |
|  | Sens (%) | Spec (%) | Acc (%) | AUC | MCC | Sens (%) | Spec (%) | Acc (%) | AUC | MCC |
| Text CNN | 96.00 | 93.00 | 94.00 | 0.98 | 0.88 | 87.00 | 79.00 | 83.00 | 0.90 | 0.67 |
| TabNet | 79.00 | 80.00 | 8.00 | 0.81 | 0.61 | 73.00 | 75.00 | 74.00 | 0.75 | 0.57 |

The motifs exclusively seen in peptide hormones have been found using the MERCI software. The list of motifs found in hormones peptides is mentioned in the Table 3. A total of 12 motifs have been identified in hormones with a coverage of 59 hormone sequences. It was observed that amino acids like phenylalanine (F), glycine (G), leucine (L), proline (P), arginine (R), tryptophan (W), and methionine (M) are the most recurring amino acids in the motifs of hormonal peptide sequences.

**Table 3:** Motif coverage of peptide hormones

| Motifs in hormone sequences | No of hormone sequences | Motifs in hormone sequences | No of hormone sequences |
| --- | --- | --- | --- |
| F G P R | 32 | L C G S | 27 |
| W F G P | 32 | M W F | 25 |
| W F G P R | 32 | M W F G | 25 |
| F G P R L | 30 | M W F G P | 25 |
| G P R L | 30 | M W F G P R | 25 |
| W F G P R L | 30 | M W F G P R L | 25 |

### BLAST performance

In this study, we performed a BLAST search to identify hormonal sequences based on similarity between the two sequences. To perform this algorithm, we used BLAST shortp which is used for the small protein sequences, i.e., peptides. We used the standard top hit approach on various e-values. The results of BLAST for evalues varying from 10^−6^ to 10^2^ are explained in Table 4.

**Table 4.**
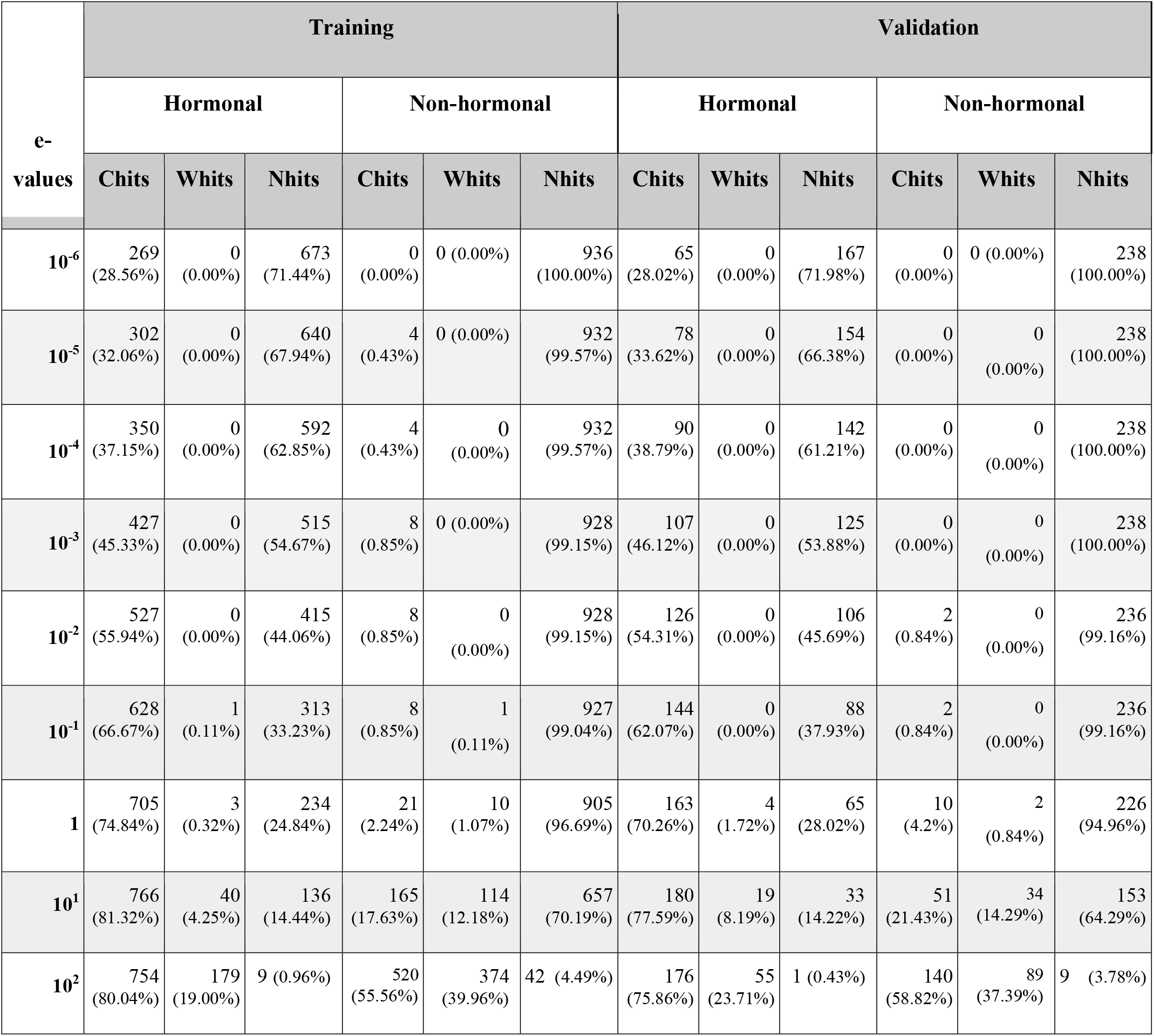
BLAST-based search results for training and validation dataset (here, Whits = wrong hits, Chits = correct hits, Nhits = no hits)

| e-values | Training |  |  |  |  |  | Validation |  |  |  |  |  |
| --- | --- | --- | --- | --- | --- | --- | --- | --- | --- | --- | --- | --- |
|  | Hormonal |  |  | Non-hormonal |  |  | Hormonal |  |  | Non-hormonal |  |  |
|  | Chits | Whits | Nhits | Chits | Whits | Nhits | Chits | Whits | Nhits | Chits | Whits | Nhits |
| $10^{-6}$ | 269<br>(28.56%) | 0<br>(0.00%) | 673<br>(71.44%) | 0<br>(0.00%) | 0 (0.00%) | 936<br>(100.00%) | 65<br>(28.02%) | 0<br>(0.00%) | 167<br>(71.98%) | 0<br>(0.00%) | 0 (0.00%) | 238<br>(100.00%) |
| $10^{-5}$ | 302<br>(32.06%) | 0<br>(0.00%) | 640<br>(67.94%) | 4<br>(0.43%) | 0 (0.00%) | 932<br>(99.57%) | 78<br>(33.62%) | 0<br>(0.00%) | 154<br>(66.38%) | 0<br>(0.00%) | 0<br>(0.00%) | 238<br>(100.00%) |
| $10^{-4}$ | 350<br>(37.15%) | 0<br>(0.00%) | 592<br>(62.85%) | 4<br>(0.43%) | 0<br>(0.00%) | 932<br>(99.57%) | 90<br>(38.79%) | 0<br>(0.00%) | 142<br>(61.21%) | 0<br>(0.00%) | 0<br>(0.00%) | 238<br>(100.00%) |
| $10^{-3}$ | 427<br>(45.33%) | 0<br>(0.00%) | 515<br>(54.67%) | 8<br>(0.85%) | 0 (0.00%) | 928<br>(99.15%) | 107<br>(46.12%) | 0<br>(0.00%) | 125<br>(53.88%) | 0<br>(0.00%) | 0<br>(0.00%) | 238<br>(100.00%) |
| $10^{-2}$ | 527<br>(55.94%) | 0<br>(0.00%) | 415<br>(44.06%) | 8<br>(0.85%) | 0<br>(0.00%) | 928<br>(99.15%) | 126<br>(54.31%) | 0<br>(0.00%) | 106<br>(45.69%) | 2<br>(0.84%) | 0<br>(0.00%) | 236<br>(99.16%) |
| $10^{-1}$ | 628<br>(66.67%) | 1<br>(0.11%) | 313<br>(33.23%) | 8<br>(0.85%) | 1<br>(0.11%) | 927<br>(99.04%) | 144<br>(62.07%) | 0<br>(0.00%) | 88<br>(37.93%) | 2<br>(0.84%) | 0<br>(0.00%) | 236<br>(99.16%) |
| **1** | 705<br>(74.84%) | 3<br>(0.32%) | 234<br>(24.84%) | 21<br>(2.24%) | 10<br>(1.07%) | 905<br>(96.69%) | 163<br>(70.26%) | 4<br>(1.72%) | 65<br>(28.02%) | 10<br>(4.2%) | 2<br>(0.84%) | 226<br>(94.96%) |
| $10^1$ | 766<br>(81.32%) | 40<br>(4.25%) | 136<br>(14.44%) | 165<br>(17.63%) | 114<br>(12.18%) | 657<br>(70.19%) | 180<br>(77.59%) | 19<br>(8.19%) | 33<br>(14.22%) | 51<br>(21.43%) | 34<br>(14.29%) | 153<br>(64.29%) |
| $10^2$ | 754<br>(80.04%) | 179<br>(19.00%) | 9 (0.96%) | 520<br>(55.56%) | 374<br>(39.96%) | 42 (4.49%) | 176<br>(75.86%) | 55<br>(23.71%) | 1 (0.43%) | 140<br>(58.82%) | 89<br>(37.39%) | 9 (3.78%) |

### Hybrid approach

In the end, various strategies were combined to get around the shortcomings of distinct approaches. These methods were created to more accurately detect the peptide hormones. This combines BLAST and motif-based methods with machine learning models. At first, peptides were categorized using the motif-search method followed by BLAST at various e-values. Not all peptides were categorized using these methods. Hence ML approach was also used in combination. The coverage and precision were much improved by the hybrid approach, which was not possible when using each of these approaches separately.

Firstly, we combined the motif-search approach with ML models to boost the performance of the classification model. The best performing algorithm – Logistic Regression (LR) achieves an AUROC of 0.95 on the training dataset and 0.94 on the validation dataset. An MCC of 0.76 and 0.74 were achieved for training and validation sets respectively.

Secondly, we combined both BLAST for e-value = 10^−1^ and motif-search with the ML model to boost the performance of our classification model. The ML model when combined with BLAST at e-value = 10^−1^ was giving the best results for all ML algorithms as compared to other e-values. The performance of the LR-based model increases from AUC 0.95 to 0.97 and 0.94 to 0.96 on the training and validation datasets for the performance when we combined BLAST with the previous model (ML + Motif-search +BLAST). The combined performance for both hybrid approaches (ML + Motif-search) and (ML + Motif-search + BLAST) is displayed in Table 5 and the AUROC plots comparing only ML model and our best performing hybrid model (ML + Motif-search + BLAST) is given in Figure 4.

**Table 5:** The performance of ML-based models in combination with Motif-search and BLAST (e-value = 10^−1^)

| ML + Motif-search |  |  |  |  |  |  |  |  |  |  |
| --- | --- | --- | --- | --- | --- | --- | --- | --- | --- | --- |
| Model | Training |  |  |  |  | Validation |  |  |  |  |
|  | Sens (%) | Spec (%) | Acc (%) | AUC | MCC | Sens (%) | Spec (%) | Acc (%) | AUC | MCC |
| **RF** | 84.5 | 83.55 | 84.03 | 0.92 | 0.68 | 81.03 | 86.13 | 83.62 | 0.92 | 0.67 |
| **LR** | **87.90** | **87.93** | **87.91** | **0.95** | **0.76** | **86.64** | **87.39** | **87.02** | **0.94** | **0.74** |
| **XGB** | 85.03 | 84.51 | 84.77 | 0.92 | 0.70 | 80.17 | 85.29 | 82.77 | 0.91 | 0.66 |
| **KNN** | 58.39 | 52.46 | 55.43 | 0.59 | 0.11 | 58.19 | 57.56 | 57.87 | 0.67 | 0.16 |
| **GNB** | 26.86 | 96.05 | 61.34 | 0.78 | 0.32 | 25.43 | 97.9 | 62.13 | 0.85 | 0.34 |
| **DT** | 73.78 | 73.82 | 73.8 | 0.80 | 0.48 | 64.22 | 75.21 | 69.79 | 0.76 | 0.40 |
| ML + Motif-search + BLAST |  |  |  |  |  |  |  |  |  |  |
|  | Training |  |  |  |  | Validation |  |  |  |  |
|  | Sens (%) | Spec (%) | Acc (%) | AUC | MCC | Sens (%) | Spec (%) | Acc (%) | AUC | MCC |

| Model | Sens (%) | Spec (%) | Acc (%) | AUC | MCC | Sens (%) | Spec (%) | Acc (%) | AUC | MCC |
| --- | --- | --- | --- | --- | --- | --- | --- | --- | --- | --- |
| **RF** | 88.96 | 85.90 | 87.43 | 0.95 | 0.75 | 86.21 | 88.66 | 87.45 | 0.95 | 0.75 |
| **LR** | **91.19** | **90.38** | **90.79** | **0.97** | **0.82** | **90.09** | **89.5** | **89.79** | **0.96** | **0.80** |
| **XGB** | 88.96 | 87.71 | 88.71 | 0.96 | 0.77 | 85.34 | 87.82 | 86.6 | 0.95 | 0.73 |
| **KNN** | 77.18 | 72.54 | 73.87 | 0.85 | 0.50 | 74.14 | 75.63 | 74.89 | 0.86 | 0.50 |
| **GNB** | 69.43 | 96.26 | 82.80 | 0.90 | 0.68 | 64.66 | 97.9 | 81.49 | 0.91 | 0.67 |
| **DT** | 81.95 | 79.70 | 80.83 | 0.89 | 0.62 | 77.16 | 76.89 | 77.02 | 0.86 | 0.54 |

**Figure 4:**
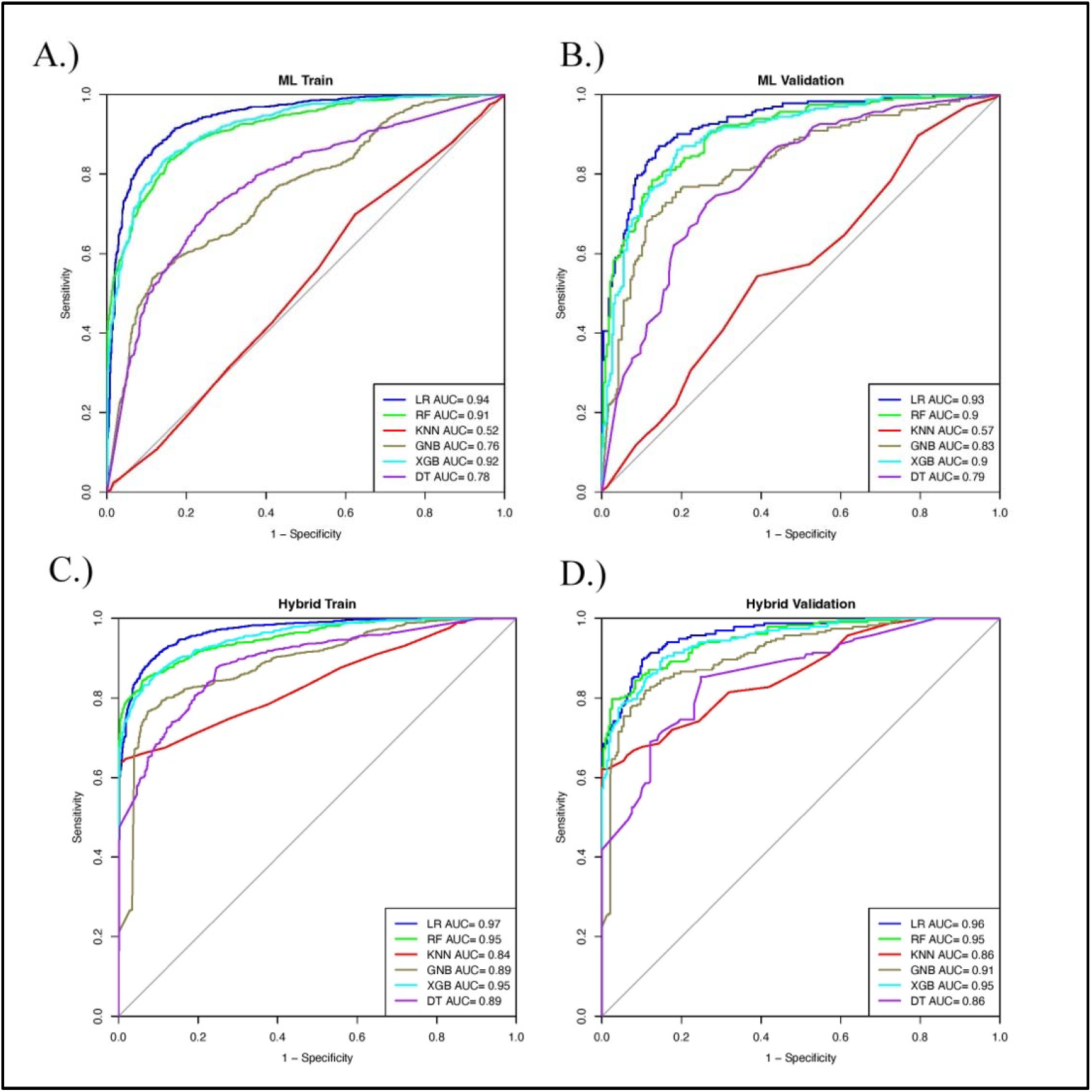
AUC plots for A) Training set in the ML model, B) Validation set in the ML model, C) Training set in the hybrid model (ML + Motif-search +BLAST) and D) Validation set in the hybrid model (ML + Motif-search +BLAST)

### Webserver and Standalone Software

The webserver for HOPPred (https://webs.iiitd.edu.in/raghava/hoppred/) that we created can distinguish between peptide hormones and non-hormone peptides. The front end and back end of the web server were created using HTML5, Java, CSS3, and PHP scripts. All of the most recent gadgets, including smartphones, tablets, iMacs, and desktop PCs, are all compatible with this web server. Predict, design, and peptide design modules are the primary modules on HOPPred.

## Discussions

Over the past few decades, peptide therapeutics have attracted much attention. The rise in publications and the creation of in silico tools and databases are evidence of this. Several peptide-based medications have previously received FDA approval due to their benefits over conventional small molecule-based medications. Peptide hormones have demonstrated promising results in replacement therapies since the very beginning [3]. There is a need for in-silico techniques that can predict peptide hormones with high confidence because the identification and screening of putative peptide hormones as drugs in the wet lab is a time-consuming, expensive, and labour-intensive operation.

We created a novel model for the current study using 1174 peptide hormones and 1174 non-hormone peptides for the prediction of peptide hormones. We have used multiple approaches to classify hormonal and non-hormonal peptides – a) motif-search, b) BLAST search, c) Machine Learning models, d) Deep Learning models, and e) hybrid models. In motif-search, we found motifs that were exclusively present in hormonal sequences. However, this approach only covered 59 hormonal sequences in total. It was found that phenylalanine (F), glycine (G), leucine (L), proline (P), arginine (R), tryptophan (W), and methionine (M) amino acids recurred in the 12 motifs found in hormonal sequences. In BLAST-search, we used the top-hit method on various e-values ranging from 10^−6^ to 10^2^. Similar to the motif approach, BLAST was not able to cover all sequences.

In ML models, we extracted 9149 compositional features using Pfeature software and selected the top performing features to develop our model. The ML-model alone was able to achieve an AUROC of 0.89, 0.90, and 0.93 for independent validation set on AAC, top 30 features and top 50 features respectively. For DL models – TextCNN and TabNet, the performance of AUROC was 0.90 and 0.75 respectively for independent validation set. As it was observed that top 50 features were giving the best performance on logistic regression (ML model). The top 50 compositional features included features like amino acid composition (AAC), dipeptide composition (DPC), tripeptide composition (TPC), physicochemical properties (PCP), distance distribution (DDR), shannon entropy (SEP), conjoint triad descriptors (CTC), composition enhanced transition and distribution (CeTD), and quasi-sequence order (QSO). We decided to combine the best performing ML model with other approaches used in the study and called it a hybrid model. Firstly, we combined the ML model with motif-search which achieved an AUROC of 0.94 on validation set. Secondly, we merged our best ML model with both motif-search and BLAST-search (e-value = 10^−1^) which achieved the highest AUROC of 0.96 with an accuracy of 89.70%, specificity of 90.09% and sensitivity of 89.50%. In addition, we performed amino acid composition analysis on the sequences to identify which amino acids are more or less likely to be found in hormones. It was observed that cysteine, aspartic acid, phenylalanine, glycine, arginine, serine, asparagine, proline, and tyrosine were significantly increased in hormonal peptide sequences whereas the compositions of amino acids like glutamic acid, isoleucine, leucine, methionine, glutamine, lysine, threonine, and valine were significantly decreased in hormonal peptide sequences than non-hormonal peptide sequences.

We have employed our best performing hybrid model on the webserver of HOPPred. We have developed a comprehensive platform that enables users to categorize both hormone and non-hormone peptide sequences. We anticipate that the scientific community engaged in peptide therapeutic research will find our study to be helpful. To assist the scientific community and promote the use of this prediction method in research, the webpage is publicly accessible. We have used the hybrid model, which is our best-performing model, in this web server. We anticipate that our prediction technique will be used by researchers around the world to create more accurate and efficient peptide-based treatment approaches for a variety of diseases.

## Author’s contributions

DK and AA collected and processed the data. DK, PV, and AA implemented the algorithms. DK, PV, and AA developed the prediction models. DK and AA developed the front end and back end of the web server. DK, AA, and GPSR prepared the manuscript. GPSR conceived and coordinated the project. All authors have read and approved the final manuscript.

## Acknowledgements

Authors are thankful to Council of Scientific and Industrial Research (CSIR) and All India Council for Technical Education for providing fellowships and the financial support. Authors are also thankful to the Department of Computational Biology, IIITD, New Delhi for infrastructure and facilities. We thank the Department of Biotechnology (DBT) for providing an infrastructure grant to the institute. We would like to acknowledge that figures were created using BioRender.com.

## Data Availability Statement

All the datasets generated in this study are available at https://webs.iiitd.edu.in/raghava/hoppred/dataset.php.

## References

1. Kołodziejski PA, Pruszyńska-Oszmałek E, Wojciechowicz T, et al. The Role of Peptide Hormones Discovered in the 21st Century in the Regulation of Adipose Tissue Functions. Genes (Basel) 2021; 12

2. Thakur SS. Proteomics and its application in endocrine disorders. Biochim Biophys Acta Proteins Proteom 2021; 1869:140701

3. Wang L, Wang N, Zhang W, et al. Therapeutic peptides: current applications and future directions. Signal Transduct Target Ther 2022; 7:48

4. Lau JL, Dunn MK. Therapeutic peptides: Historical perspectives, current development trends, and future directions. Bioorg Med Chem 2018; 26:2700–2707

5. Yan K, Lv H, Guo Y, et al. TPpred-ATMV: therapeutic peptide prediction by adaptive multiview tensor learning model. Bioinformatics 2022; 38:2712–2718

6. Shoombuatong W, Schaduangrat N, Pratiwi R, et al. THPep: A machine learning-based approach for predicting tumor homing peptides. Comput Biol Chem 2019; 80:441–451

7. Agrawal P, Bhagat D, Mahalwal M, et al. AntiCP 2.0: an updated model for predicting anticancer peptides. Brief Bioinform 2021; 22

8. Yan W, Tang W, Wang L, et al. PrMFTP: Multi-functional therapeutic peptides prediction based on multi-head self-attention mechanism and class weight optimization. PLoS Comput Biol 2022; 18:e1010511

9. Kaur D, Arora A, Patiyal S, et al. Hmrbase2: A comprehensive database of hormones and their receptors.

10. Fu L, Niu B, Zhu Z, et al. CD-HIT: accelerated for clustering the next-generation sequencing data. Bioinformatics 2012; 28:3150–2

11. Deutsch EW, Lam H, Aebersold R. PeptideAtlas: a resource for target selection for emerging targeted proteomics workflows. EMBO Rep 2008; 9:429–34

12. Pande A, Patiyal S, Lathwal A, et al. Pfeature: A Tool for Computing Wide Range of Protein Features and Building Prediction Models. J Comput Biol 2023; 30:204–222

13. Ren K, Zeng Y, Cao Z, et al. ID-RDRL: a deep reinforcement learning-based feature selection intrusion detection model. Sci Rep 2022; 12:15370

14. Abraham A, Pedregosa F, Eickenberg M, et al. Machine learning for neuroimaging with scikit-learn. Front Neuroinform 2014; 8:14

15. Di Noto T, von Spiczak J, Mannil M, et al. Radiomics for Distinguishing Myocardial Infarction from Myocarditis at Late Gadolinium Enhancement at MRI: Comparison with Subjective Visual Analysis. Radiol Cardiothorac Imaging 2019; 1:e180026

16. Flayer CH, Perner C, Sokol CL. A decision tree model for neuroimmune guidance of allergic immunity. Immunol Cell Biol 2021; 99:936–948

17. Yi Y, Sun D, Li P, et al. Unsupervised random forest for affinity estimation. Comput Vis Media (Beijing) 2022; 8:257–272

18. Stoltzfus JC. Logistic regression: a brief primer. Acad Emerg Med 2011; 18:1099–104

19. Miao Y, Hunter A, Georgilas I. An Occupancy Mapping Method Based on K-Nearest Neighbours. Sensors (Basel) 2021; 22

20. Joshi D, Mishra A, Anand S. A naïve Gaussian Bayes classifier for detection of mental activity in gait signature. Comput Methods Biomech Biomed Engin 2012; 15:411–6

21. Hou N, Li M, He L, et al. Predicting 30-days mortality for MIMIC-III patients with sepsis-3: a machine learning approach using XGboost. J Transl Med 2020; 18:462

22. Sercan S, Arık S, Pfister T. TabNet: Attentive Interpretable Tabular Learning. 2021

23. Kalchbrenner N, Grefenstette E, Blunsom P. A Convolutional Neural Network for Modelling Sentences. 2014;

24. Sharma N, Patiyal S, Dhall A, et al. AlgPred 2.0: an improved method for predicting allergenic proteins and mapping of IgE epitopes. Brief Bioinform 2021; 22:

25. Altschul SF, Gish W, Miller W, et al. Basic local alignment search tool. J Mol Biol 1990; 215:403–10

26. Arora A, Patiyal S, Sharma N, et al. A random forest model for predicting exosomal proteins using evolutionary information and motifs.

27. Vens C, Rosso M-N, Danchin EGJ. Identifying discriminative classification-based motifs in biological sequences. Bioinformatics 2011; 27:1231–8

