## Supplementary Table S1 for "Prediction of peptide hormones using an ensemble of machine learning and similarity-based methods"

Top 50 Features used for machine learning techniques

| Feature | Description |
|---|---|
| DPC1_CF | Dipeptide composition of Cysteine-Phenylalanine |
| TPC_AQI | Tripeptide Composition of Alanine Glutamine Isoleucine |
| TPC_CFN | Tripeptide Composition of Cysteine Proline henylalanine Asparagine |
| TPC_CVL | Tripeptide Composition of Cysteine Valine Leucine |
| TPC_DLN | Tripeptide Composition of Aspartic acid Leucine Asparagine |
| TPC_DML | Tripeptide Composition of Aspartic acid Methionine Leucine |
| TPC_EQS | Tripeptide Composition of Glycine lutamic acid Glutamine Serine |
| TPC_FRP | Tripeptide Composition of Proline henylalanine Arginine Proline |
| TPC_GCK | Tripeptide Composition of Glycine Cysteine Lysine |
| TPC_GLM | Tripeptide Composition of Glycine Leucine Methionine |
| TPC_GNF | Tripeptide Composition of Glycine Asparagine Proline henylalanine |
| TPC_HLC | Tripeptide Composition of Histidine Leucine Cysteine |
| TPC_KCC | Tripeptide Composition of Lysine Cysteine Cysteine |
| TPC_KYS | Tripeptide Composition of Lysine Tyrosine Serine |
| TPC_LAN | Tripeptide Composition of Leucine Alanine Asparagine |
| TPC_LGM | Tripeptide Composition of Leucine Glycine Methionine |
| TPC_LLF | Tripeptide Composition of Leucine Leucine Proline henylalanine |
| TPC_LMG | Tripeptide Composition of Leucine Methionine Glycine |
| TPC_LNS | Tripeptide Composition of Leucine Asparagine Serine |
| TPC_MAY | Tripeptide Composition of Methionine Alanine Tyrosine |
| TPC_NTP | Tripeptide Composition of Asparagine Threonine Proline |
| TPC_RGL | Tripeptide Composition of Arginine Glycine Leucine |
| TPC_RKY | Tripeptide Composition of Arginine Lysine Tyrosine |
| TPC_RRP | Tripeptide Composition of Arginine Arginine Proline |
| TPC_SIR | Tripeptide Composition of Serine Isoleucine Arginine |
| TPC_THR | Tripeptide Composition of Threonine Histidine Arginine |
| TPC_TTG | Tripeptide Composition of Threonine Threonine Glycine |
| TPC_VCG | Tripeptide Composition of Valine Cysteine Glycine |
| TPC_VSF | Tripeptide Composition of Valine Serine Proline henylalanine |
| PCP_AR | Composition of aromatic residues |
| DDR_C | Distance distribution of Cysteine |
| DDR_K | Distance distribution of Lysine |
| SER_T | Shannon entropy of Threonine |

| | |
|---|---|
| SEP_LR | Shannon entropy of large residues |
| CTC_174 | Conjoint Triad Descriptors, Normalize frequency of Group 1 (A, G, V)-Group 6 (C)-Group 4 (H, N, Q, W) |
| CTC_374 | Conjoint Triad Descriptors, Normalize frequency of Group 3 (Y, M, T, S)-Group 6 (C)-Group 4 (H, N, Q, W) |
| CTC_477 | Conjoint Triad Descriptors, Normalize frequency of Group 4 (H, N, Q, W)-Group 6 (C)-Group 6 (C) |
| CTC_677 | Conjoint Triad Descriptors, Normalize frequency of Group 6 (D, E)-Group 6 (C)-Group 6 (C) |
| CTC_713 | Conjoint Triad Descriptors, Normalize frequency of Group 6 (C)-Group 1 (A, G, V)-Group 3 (Y, M, T, S) |
| CeTD_11_PO | Number of transitions takes place from L,I,F,W,C,M,V,Y residues to L,I,F,W,C,M,V,Y residues for polarity attribute |
| CeTD_11_SS | Number of transitions takes place from E,A,L,M,Q,K,R,H residues to E,A,L,M,Q,K,R,H residues for secondary structure attribute |
| CeTD_11_SA | Number of transitions takes place from A,L,F,C,G,I,V,W residues to A,L,F,C,G,I,V,W residues for solvent accessibility attribute |
| CeTD_12_HB | Number of transitions takes place from R,K,E,D,Q,N residues to G,A,S,T,P,H,Y residues for hydrophobicity attribute |
| CeTD_13_SA | Number of transitions takes place from A,L,F,C,G,I,V,W residues to M,S P,T,H,Y residues for solvent accessibility attribute |
| CeTD_21_HB | Number of transitions takes place from G,A,S,T,P,H,Y residues to R,K,E,D,Q,N residuesvfor hydrophobicity attribute |
| CeTD_21_PZ | Number of transitions takes place from C,P,N,V,E,Q,I,L residues to G,A,S,D,T residues for polarizability attribute |
| CeTD_23_PO | Number of transitions takes place from P,A,T,G,S residues to H,Q,R,K,N,E,D residues for polarity attribute |
| CeTD_100_p_PO1 | Number of L,I,F,W,C,M,V,Y residues for polarity present in 100% quartile |
| CeTD_100_p_PZ3 | Number of K,M,H,F,R,Y,W residues for polarizability attribute present in 100% quartile |
| QSO1_SC_V | Quasi-sequence order with Schneider matrix for Valine |
